# Validation of differentiated sinoatrial-like hiPSCs as a model of native sinus node myocytes

**DOI:** 10.64898/2026.01.21.700785

**Authors:** Eleonora Torre, Yvonne Sleiman, Haikel Dridi, Azzouz Charrabi, Nadia Mekrane, Giuseppe Angelini, Leïla Talssi, Rajesh K. Soni, Valentina Di Biase, Heloise Faure-Gautron, Pascal Seyer, Pieter de Tombe, Andrew R. Marks, Jean-Luc Pasquié, Alain Lacampagne, Matteo E. Mangoni, Pietro Mesirca, Albano C. Meli

## Abstract

**Background:** Human induced pluripotent stem cell derived cardiomyocytes (hiPSC-CMs) constitute an attractive system for basic research and pharmacologic screening of new molecules of clinical interest. Numerous protocols aiming at differentiating atrial- or ventricular-like cardiomyocytes (hiPSC-CMs) are available. Conversely, only a few are available for obtaining patient-derived sinoatrial node-like pacemaker myocytes (PM-hiPSC-CMs).

Here we validate a new protocol to differentiate mature PM-hiPSC-CMs as a model of native sinoatrial node (SAN) myocytes.

**Methods:** We generated PM-hiPSC-CMs through a 2D matrix-sandwich method promoting epithelial-to-mesenchymal transition and small molecule-based temporal modulation of Wnt signaling pathway. In addition, we treated our cells with triiodothyronine, dexamethasone and intracellular cyclic AMP (DTA) to enhance expression of proteins involved in intracellular Ca^2+^ handling.

**Results:** Proteomic analyses showed expression of key SAN proteins in DTA-treated PM-hiPSC-CMs. Importantly, expression of proteins related to Ca^2+^ handling was increased in DTA-treated PM-hiPSC-CMs compared to untreated ones. DTA-treated PM-hiPSC-CMs displayed action potentials, ionic currents and intracellular Ca^2+^ dynamics typical of native SAN. In addition, pacemaker activity responded to both β-adrenergic and muscarinic stimulation.

**Conclusions:** Our data indicate that the differentiation protocol effectively generates PM-hiPSC-CMs with typical native human SAN features. This protocol may serve as a potential approach to generate PM-hiPSC-CMs from patients with history of sinoatrial node disfunction (SND) carrying different mutations in ion channels underlying pacemaking. In addition, these *in vitro* models of SND could be used for testing long-term vector-based gene therapeutic strategies to handle bradycardia.

## INTRODUCTION

Human induced pluripotent stem cells (hiPSCs) are able to differentiate into multiple cell lineages, thereby constituting convenient models of a wide range of human diseases. hiPSCs can be generated using skin biopsies, urine or blood samples from affected individuals. Techniques have been developed for generating hiPSC from human tissues ^1,2^ and more recently, hiPSC-derived cardiomyocytes (hiPSC-CMs) were used to investigate cardiac phenotypes ^3–7^. To date, hiPSC-CMs have been mostly limited to modelling of inherited cardiopathies affecting ventricular cardiomyocytes ^6,7^. In addition, the majority of cardiac differentiation protocols using hiPSC yielded mixed populations of cardiomyocytes composed primarily of ventricular cells (ventricular-like hiPSC-CMs) containing limited number of atrial-and pacemaker-like cells. Hence, the current knowledge in the field is mainly focused on differentiated ventricular-like hiPSC-CMs.

The generation of functional and mature sinoatrial node (SAN) -like pacemaker myocytes (PM-hiPSC-CMs) is an important step for opening the way to modeling atrial and nodal inherited arrhythmias such as primary sinus node dysfunction (SND) in patient-specific context. Treatment of SND only includes a few options and the vast majority of patients with irreversible SND symptoms can only be treated with permanent pacemaker implantation, which stands as the primary and definitive therapy ^8^. The development of an *in vitro* model of inherited SND using patient-derived cardiomyocytes would be of paramount importance to understand further the mechanisms underlying SND and test innovative clinical strategies. In this context, few protocols have obtained PM-hiPSC-CMs (for review see ^9,10^) using small-molecule protocols ^11–14^ or co-culture with visceral endoderm-like cell line leading to mixed cell population ^15^.

Evidence suggest significant roles of triiodothyronine (T3), dexamethasone (Dex) and intracellular cyclic AMP (cAMP) to improve key properties of hiPSC-CMs ^16–18^. Since Dex accelerates Ca^2+^ transient decay enhancing sarcoplasmic reticulum (SR) Ca^2+^ ATPase (SERCA) and Na^+^-Ca^2+^ exchanger (NCX) function in human embryonic stem cell derived ventricular-like cardiomyocytes (hESC-CMs) ^19^, it is considered a good candidate molecule to promote hiPSC-CM maturation when used in combination with other factors such as T3. Notably, association of T3 and Dex ameliorates the functional coupling between L-type Ca^2+^ channels and ryanodine receptor (RyR2) on SR in hiPSC-CMs, enhancing the Ca^2+^-induced Ca^2+^-release (CICR) ^17^. Furthermore, persistent activation of the cAMP pathway contributes to hiPSC-CM maturation by positively affecting the assembly of gap junctions ^16^. Finally, cell behavior and stem cell fate choices are also critically impacted by extracellular matrix ^20^. Sandwiching hiPSCs by overlaying the 2D cardiac sheet with Matrigel promoted an epithelial-to-mesenchymal transition, which is an essential first step in generating cardiac progenitors. Furthermore, combining the Matrigel overlay with application of growth factors results in the generation of cardiomyocytes in high yield and purity from multiple hiPSCs lines ^21^.

Here, we aim to characterize PM-hiPSC-CMs obtained using unique 2D-sheets - matrix-sandwich method-based - ^21^ cardiac differentiation protocol combined with Dex, T3 and intracellular cAMP (DTA) treatments ^16–18^ allowed to provide a robust *in vitro* model for studying SAN pathophysiology.

## METHODS

### Cell line

Male hiPSC cell line AG08C5 was obtained from a healthy donor (from NeuroMyoGène Institute, Lyon) as previously described ^5,22^. The cell line was maintained in standard conditions (21% O_2_, 5% CO_2_ and 37°C) on hES (human embryonic stem cell)-qualified Matrigel (Corning). hiPSC cells were enzymatically dissociated using TrypLE enzyme (Gibco) and passaged every 4 days.

### 2D cardiac sheet differentiation

2D-sandwich based protocol was used to differentiate hiPSC line to PM-hiPSC-CMs (Fig. 1A). Briefly, undifferentiated single hiPSCs were plated into 6 well dishes in standard conditions (21% O_2_ and 5% CO_2_ at 37 ° C). At 90% confluency (day-1), 0.04 mg of Matrigel reduced growth factor (MgFr, Corning) was added in Stemflex medium (Gibco). On day 0, mesoderm was induced by adding 10µM CHIR99021 and 0.04 mg of MgFr in glucose rich RPMI 1640-B27-minus insulin medium (Gibco). On day 1, the medium was changed with RPMI 1640-B27-minus insulin only. On day 3, differentiation of cardiac progenitor was induced by adding 5 µM Wnt Antagonist II (IWP2, Sigma) in the RPMI 1640-B27-minus insulin medium; the medium was kept 2 days. On day 5, according to Liu et al., 2020, the medium was changed and 1.25 ng/ml human Bone Morphogenetic Protein 4 (BMP4, Sigma), 960 nM PD173074 (FGFR1 inhibitor, Tocris) and 1 µM BMS189453 (Retinoic Acid inhibitor, Sigma) were added in RPMI 1640-B27-minus insulin medium. On day 7, the medium was changed to RPMI 1640-B27 (Gibco) and was kept until day 32; the medium was renewed every 2 days. On day 32, the cells were treated for 8 days with 1 µM Dexamethasone (Sigma), 100 nM T3 (Sigma) and 500 µM dibutyryl cAMP (Sigma), here referred to as DTA treatment, to improve phenotype maturity, as previously proposed ^16,17^. The medium was renewed every 2 days. DTA-treated and untreated PM-hiPSC-CMs were maintained in culture until day 40-50 of differentiation.

**Fig 1.**
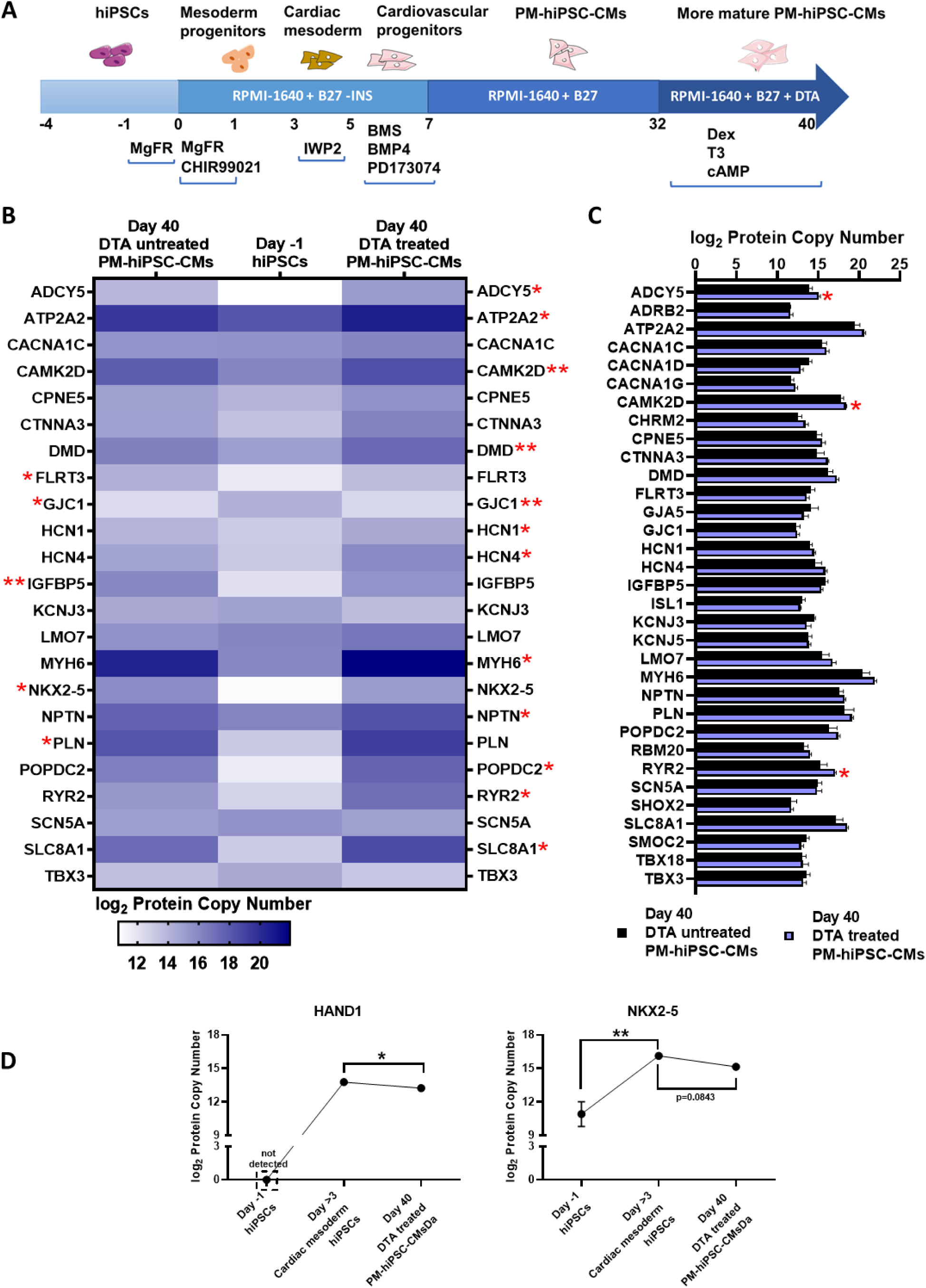
Differentiation protocol and proteomic profiles of PM-hiPSC-CMs. (A) Schematic diagram of PM-hiPSC-CMs differentiation protocol with DTA treatment. (B) Heatmap displaying log_2_ mean value of copy number of 23 different proteins measured by global quantitative proteomic at day-1 and day 40 of differentiation protocol to obtain DTA-treated and untreated PM-hiPSC-CMs. d≥3 per condition. Statistic: unpaired non-parametric Kruskal-Wallis test followed by Dunn’s multiple comparison test. *, *p* < 0.05. **, *p* < 0.01 vs Day-1 hiPSCs. (C) Histograms of log_2_ mean value of copy number of 33 different proteins (including proteins not expressed at day -1) measured by global quantitative proteomic in DTA-treated and untreated PM-hiPSC-CMs at day 40 of differentiation. d≥3 per condition. Statistic: unpaired non-parametric Mann-Whitney test. *, *p* < 0.05 vs untreated PM-hiPSC-CMs. (D) Variation of log_2_ mean value of copy number of HAND1 and NKX2-5 proteins during differentiation protocol (Day -1 hiPSCs, cardiac mesoderm and DTA-treated PM-hiPSC-CMs at day 40 of differentiation). d≥3 per condition. Statistic: unpaired non-parametric Mann-Whitney test for HAND1 and unpaired non-parametric Kruskal-Wallis test followed by Dunn’s multiple comparison test for NKX2-5. *, *p* < 0.05. **, *p* < 0.01.

### 2D cardiac sheet dissociation of PM-hiPSC-CMs

Beating 2D cardiac sheets were dissociated using TrypLE enzyme to obtain isolated single PM-hiPSC-CMs as previously described ^7^. Briefly, 2D cardiac sheets were washed twice with Ca^2+^-and Mg^2+^-free PBS (Sigma) and then incubated with pre-warmed TrypLE for 10min at 37°C. RPMI 1640-B27 was used to stop the enzymatic activity of TrypLE. Remaining cell clumps were filtered using a 37µm reversible strainer (Stemcell Technologies). Whole lysate was then centrifuged for 5min at 200g and cell pellet resuspended with RPMI 1640-B27 containing ROCK inhibitor (Miltenyi Biotec).

### Global quantitative proteomic analysis

For global quantitative proteomics of PM-hiPSC-CMs (treated with, +DTA, or not, -DTA), diaPASEF (data independent acquisition) based proteomic was performed with a minimum of three biological repetitions per group. The cut-off values for differentially expressed proteins included p-value<0.05 (permutation-based FDR correction), |fold-change| ≥1.5, and unique peptides ≥2.

### Gene Ontology analysis

Gene Ontology Analysis was performed on differentially expressed proteins (DEPs) in +DTA versus -DTA groups identified by 3 different proteomic quantifications. Protein names were converted into “Entrez ID” using the “org.Hs.eg.db” package (version 3.21.0) ^23^. Up- and down-regulated DEPs were separately analyzed evaluating the GO terms associated to the Biological Process (BP) category. Enriched BP terms were obtained exploiting the “enrichGO” function of the “clusterProfiler” package (v4.16.0) ^24,25^ in R software (v4.5.1)^26^ and selecting only the statistically significant terms based on the following parameters: fold enrichment = GeneRatio / Background Ratio; p-value<0.05; q-value<0.05; Benjamini-Hochberg correction.

Among the most significant terms, BP terms were selected according to their biological relevance, removing generic or redundant terms. The treeplot of the selected BP terms was generated by the “treeplot” function in the “clusterProfiler” package using the “pairwise_termism” function to calculate the semantic similarity based on median. The treeplot graphic style was modified by the “ggplot” function in the “ggtree” package (v3.16.3)^27^.

### 2D-intracellular Ca2+ imaging using fluorescent confocal microscopy

To monitor spontaneous [Ca^2+^]i transients and evaluate sarcoplasmic-reticulum (SR) Ca^2+^ load, dissociated DTA-treated and untreated PM-hiPSC-CMs were studied using Zeiss LSM980 (AiryscanII) fluorescent confocal microscope as previously described ^28^. Briefly, PM-hiPSC-CMs were dissociated, as described above, and seeded on hES-qualified Matrigel -coated 35 mm Ø glass Fluorodish (FD3510-100, WPI) with a density of 2.5 * 10^2^ cells/mm^2^. PM-hiPSC-CMs were loaded for 20 min with the Ca^2+^ indicator 5µM Fluo4-AM dissolved in Tyrode’s solution containing (mM): 140 NaCl, 5.4 KCl, 1 MgCl_2_, 1.8 CaCl_2_, 5 HEPES, 5.5 D-Glucose; pH was set to 7.4 with NaOH. After washing Fluo4-AM twice, Ca^2+^ fluorescence in PM-hiPSC-CMs were recorded in Tyrode solution. Images were acquired by scanning the cells with an argon laser in line-scan configuration (*i.e.* x–t mode, 2.46 ms per line; 512 pixels × 10000 lines). Fluorescence was excited at 488 nm and emissions were collected at >505 nm. A 63× Plan Apo oil (1.4 NA) immersion objective was used to record intracellular Ca^2+^. Measurements were performed in perfused cells at physiological temperature (37°C) with a temperature-controlled chamber and warmed solutions. SR Ca^2+^ load was measured by acute addition of 10 mM caffeine to the bath solution. Diastolic Ca^2+^ was used as reference fluorescence (F_0_) for signal normalization (F/F_0_). Time-course of Ca^2+^ fluorescence was exported using ImageJ and Ca^2+^ transients were analyzed using Clampfit (ver. 11.1).

### Patch-clamp recording of ionic currents in PM-hiPSC-CMs

Dissociated DTA-treated PM-hiPSC-CMs were plated at density of 2.5 * 10^2^ cells/mm^2^ on 40 mm Ø polystyrene dishes (TPP, TECHNO PLASTIC PROD) pre-coated with hES-qualified Matrigel. Spontaneous action potentials and ionic currents were recorded using the whole-cell configuration, by employing an Axon 700A amplifier (Molecular Devices) coupled to a 1550B Digidata (Molecular Devices). Signals were filtered at 2 kHz via pClamp 10.4 software (Molecular Devices). Membrane capacitance (C_m_) and series resistance were measured in all cells studied. Measurements were performed in Tyrode perfused cells at physiological temperature (37 °C) with a temperature-controlled chamber and warmed solutions. To evaluate the ‘funny’ current (I_f_), 2mM BaCl_2_ was added to the Tyrode solution. 2mM Cs^+^ was added to the Tyrode solution to inhibit I_f_. Calcium currents were recorded with an extracellular solution (pH 7.4) containing (mM): 135 TEA-Cl, 2 CaCl_2_, 1 MgCl_2_, 10 HEPES, 10 4-Aminopyridine, 5.5 glucose, 0.03 tetrodotoxin (TTX). Patch pipettes (resistance 2-3 MΩ) were filled with intracellular solution (pH 7.2) containing (mM): 80 K-aspartate, 50 KCl, 5 EGTA, 2 CaCl_2_, 1 MgCl_2_, 3 ATP (Na-salt) and 5 Hepes-KOH. To record calcium-currents, intracellular solution was changed to (mM): 125 CsOH, 20 TEA-Cl, 10 HEPES, 1.2 CaCl_2_, 5 MgATP, 0.1 LiGTP, 5 EGTA (pH 7.2). I_Na_ recordings were carried out with an external solution containing (in mM): 10 NaCl, 130 TEA-Cl, 5 HEPES, 5.4 KCl, 1.8 CaCl_2_, 1 MgCl_2_, 0.005 Nifedipine (Nife), 5.5 glucose (pH 7.4). Intracellular solution was the same used for calcium current recordings. The I_f_ was elicited by applying hyperpolarizing steps in the range of -55 to -135 mV from a holding potential (HP) of -35mV in -10 mV increments. Steps duration was set to reach steady-state of activation, followed by a fully-activating step at -135 mV. I_f_ current-voltage (I-V) relationships were obtained by measuring the time-dependent current activated during the voltage steps; steady state activation curves were measured by plotting normalized tail currents measured at -135 mV^29^. Steady state activation curves were fitted by Boltzmann equation:

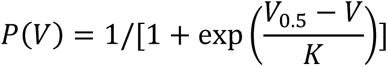

where P(V) is the voltage dependency of f-channels open probability, V_0.5_ is the half-activation voltage and k is the slope factor. Voltage-clamp (V-clamp) single step protocol (from -35 mV to -85 mV) was used to evaluate the Cs^2+^ effect on I_f_. Total Ca^2+^ current (I_CaTOT_) was elicited by applying depolarizing steps to the range of -80/+60 mV from a HP of -80 mV long enough to reach steady-state of inactivation^30^. L-type Ca^2+^ current (I_CaL_) was activated by applying depolarizing steps in the range of -55 to +60 mV from a HP of -55 mV. I_CaTOT_ and I_CaL_ I-V relationships were obtained by measuring peak current during voltage steps; T-type Ca^2+^ current (I_CaT_) I-V relationship was calculated as the difference between values obtained from HP of − 55 mV from those recorded from HP − 80 mV. V-clamp single step protocol (from - 55 mV to -10 mV) was used to evaluate the dihydropyridine insensitive current component by applying 3µM Nife. Total I_Na_ was activated by applying depolarizing steps in the range of -80 to +60 mV from a HP of -55 mV. 100nM TTX was used to record TTX-resistant I_Na._ TTX-sensitive I_Na_ was obtained subtracting the TTX-resistant I_Na_ from total I_Na_. G protein gated K^+^ current (I_KACh_) was measured as 1µM Acetylcholine (ACh)-sensitive current in V-clamp gap free protocol with HP of -45 mV. Spontaneous action potentials (APs) were acquired in gap-free mode. APs were recorded before and after 1µM Isoprotenerol (Iso) or ACh superfusion. V-clamp single step protocols for I_f_ and I_CaL_ were used to evaluate the effect of 1µM Iso or ACh on DTA-treated PM-hiPSC-CMs. Current densities (pA/pF) were obtained by normalizing current amplitudes to C_m_.

### Statistical Analysis

Statistical analysis was performed using Prism 8.0 (GraphPad Software, LLC, Boston MA, USA). Normality of distribution was tested using the Shapiro-Wilk test. For general consistency, non-parametric tests were preferred over parametric tests to interpret data. Data are expressed as mean ± the standard error of the mean (SEM). p ≤ 0.05 was considered statistically significant. Statistical tests used in each experiment are specified throughout the figure legends and *P* values are indicated in panels.

## RESULTS

### Differentiation of PM-hiPSC-CMs

Based on the protocol developed by Liu et al., 2020 ^12^, we optimized the enrichment of the PM-hiPSC-CMs by adapting 2D matrix-sandwich method that further facilitates mesoderm differentiation^31^. Additionally, because PM-hiPSC-CMs display enhanced glycolytic pathway compared to the ventricular-like cells ^32–34^, glucose-rich RPMI 1640-B27 medium was used to facilitate maturation of PM-hiPSC-CMs over that of ventricular-like hiPSC-CMs (Fig 1A). Previous studies have shown the capability of thyroid and steroid hormones (*i.e.,* T3 and dexamethasone) and cellular second messengers (*i.e.,* cAMP) to improve the function and morphology of hiPSC-CMs ^16–18^. We hypothesized that a similar treatment could be applied to PM-hiPSC-CMs to promote maturity. Thus, we applied dexamethasone-T3-cAMP (DTA) treatment to differentiate PM-hiPSC-CMs (Fig 1A).

To obtain an overview of the whole protein landscape in the resulting DTA-treated and untreated PM-hiPSC-CMs, we performed global quantitative proteomic analysis (Fig 1B-D). Liu et al., 2020 ^12^ showed the effect of the modulation of Wnt pathway to drive differentiation toward SAN-like phenotype of hiPSC-CMs. Here, we assessed the effect of DTA treatment, after Wnt modulation. In particular, we investigated expression of proteins typically enriched in the SAN, as well as in the cardiac conduction system (CCS). Expression of hyperpolarisation-activated “funny” f-(HCN) channels (HCN1, HCN4) was significantly increased in DTA-treated PM-hiPSC-CMs compared to Day-1 hiPSCs, but not in DTA untreated-hiPSC-CMs (Fig 1B). Expression of connexin-45 (Cx45/GJC1) channel was increased in both DTA-treated and untreated PM-hiPSC-CMs, compared to Day-1 hiPSCs (Fig 1B). The activity of cardiac adenylate cyclase (AC) is a downstream effector of β-adrenergic receptor (β_1_- and β_2_-AR) and type 2 muscarinic receptor (M2R). Type 5 AC (ADCY5) was increased in DTA-treated PM-hiPSC-CMs at day 40 of differentiation compared to Day-1 hiPSCs (Fig 1B). It has been demonstrated that dexamethasone and T3 treatment of hiPSC-CMs induces amelioration of several functional and structural features including SR Ca^2+^ handling ^16–18^. Proteomic data showed a significant increased expression of proteins involved in intracellular SR Ca^2+^ handling (*i.e.*, ATP2A2 -SERCA2a-, CAMK2D -CaMKIIδ-, RYR2 and SLC8A -NCX1-) upon DTA treatment in PM-hiPSC-CMs at day 40 of differentiation compared to Day-1 hiPSCs, while it remained unchanged in DTA-untreated PM-hiPSC-CMs (Fig 1B). Expression of SERCA2a modulator, PLN, was significantly increased only in DTA-untreated PM-hiPSC-CMs at day 40 of differentiation compared to Day-1 hiPSCs (Fig 1B).

We found other genes expressed in PM-hiPSC-CMs at day 40 of differentiation that are enriched in the native SAN including CPNE5, IGFBP5 and SMOC2^35^. We then searched for other proteins known to be expressed in the native SAN. Previous work using Single-nucleus RNA sequencing (snRNA-seq) of native SAN architecture identified MYH6, CTNNA3, RBM20 and DMD as part of the SAN^36^. In addition, POPDC2 has recently been identified as cAMP effector proteins and have been proposed to be part of the protein network involved in cAMP signalling and function of TREK channels in the SAN^37^. Finally, LMO7 and NPTN are additional markers of the CCS^38,39^. Notably, DTA-treatment significantly increased MYH6, DMD, POPDC2 and NPTN protein levels compared to Day-1 hiPSCs (Fig 1B). IGFBP5 protein level was increased in DTA-untreated PM-hiPSC-CMs compared to Day-1 hiPSCs (Fig 1B).

In Figure 1C, we compared expression of proteins in DTA-treated and untreated PM-hiPSC-CMs at day 40 after differentiation, including proteins that were not expressed at Day-1 (*i.e.*, ADRB2, CACNA1D, CACNA1G, CHRM2, ISL1, KCNJ5, RBM20, SHOX2 and TBX18).

Results confirmed the expression of native SAN markers SHOX2, ISL1, TBX18 and TBX3 ^36,40,41^ in DTA-treated and untreated PM-hiPSC-CMs (Fig 1C), as well as expression of ion channels involved in SAN pacemaker activity: Ca^2+^ channels (Ca_v_1.2/CACNA1C, Ca_v_1.3/CACNA1D, Ca_v_3.1/CACNA1G), the cardiac isoforms of G protein activated inward rectifier K^+^ channels (GIRK1/KCNJ3, GIRK4/KCNJ5) and the cardiac TTX-resistant Na^+^ channel (Na_v_1.5/SCN5A) (Fig 1C). M2R (CHRM2) and β_2_-AR (ADRB2) were detected in DTA-treated and untreated PM-hiPSC-CMs (Fig 1C), but not the β_1_-AR (ADRB1) isoform. Among genes newly identified as markers of the SAN, we found also SMOC2 and RBM20 in DTA-treated and untreated PM-hiPSC-CMs at day 40 after differentiation. Expression of these markers was not detected at day -1 (Fig 1C).

Notably, ventricular cells are identified by expression of HAND1 and NKX2-5^9^. Expression of these markers starts to increase during cardiac mesoderm formation. In line with heart development, HAND1 was not detected at day -1 of differentiation, but at cardiac mesoderm stage (Fig 1D). HAND1 expression was significantly reduced in DTA-treated PM-hiPSC-CMs at day 40 (Fig 1D). Similarly, at day -1, NKX2-5 was present at low levels, expression was increased at the stage of cardiac mesoderm. DTA-treatment induced a trend to down-regulation of NKX2-5 in PM-hiPSC-CMs at day 40 (Fig 1D).

We performed a Gene Ontology (GO) enrichment analysis to interpret proteomic data as network of differentially expressed proteins (DEPs) involved in specific biological processes in DTA-treated versus untreated PM-hiPSC-CMs (Fig 2). GO analysis (Fig 2A) revealed that up-regulated DEPs in DTA-treated PM-hiPSC-CMs are involved in processes associated with cardiac activity, namely cardiac muscle conduction and inner calcium homeostasis, regulation of sodium ion transport, response to oxygen levels, aerobic respiration and iron ion transport regulation. Conversely, up-regulated DEPs in DTA-untreated cells (Fig 2B) were related to cell cycle progression and proliferation, vascular endothelial growth factor signalling pathway, epithelial-to-mesenchymal transition, and stress fiber assembly and were unrelated to adult cardiac function. These results confirmed that DTA treatment of hiPSC-CMs boosts the expression of proteins typically associated with heart automaticity, repressing/diminishing proteins involved in non-cardiac and proliferating functions.

**Fig 2.**
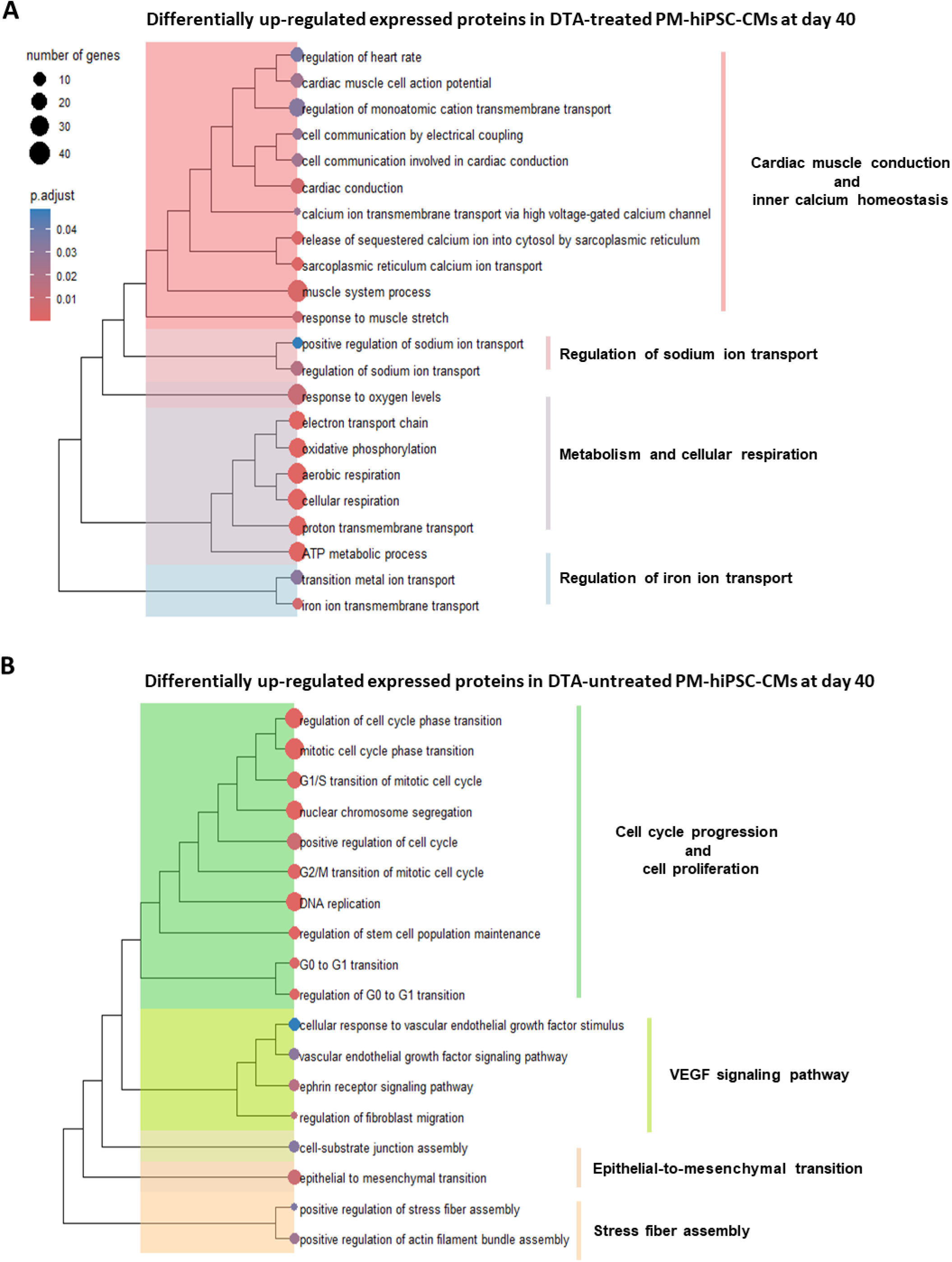
Gene Ontology (GO) analysis of proteomic analysis on DTA-treated versus untreated hiPSC-CMs. Treeplot graphs depicting the enriched GO terms of selected biological processes associated to differentially expressed proteins (DEPs) in DTA-treated (+DTA, upregulated proteins, panel A) versus untreated PM-hiPSC-CMs (-DTA, downregulated proteins, panel B) proteomic data. Cluster names of common GO terms are indicated to the right of each group. The size of each circle is related to the number of genes which are associated to the GO term; while its colour depends on the calculated adjusted p-value (p.adjust, Benjamini-Hochberg correction).

In sum, these results indicate that at day 40, DTA-treated PM-hiPSC-CMs present with reduced expression of key ventricular markers such as HAND1 and NKX2.5 and up-regulation of proteins involved in intracellular SR Ca^2+^ handling, in comparison to DTA-untreated PM-hiPSC-CMs. Taken together, these data show that DTA treatment helps drive the protein expression pattern of PM-hiPSC-CMs to a condition close to that of native SAN pacemaker myocytes.

### Intracellular Ca^2+^ handling properties of PM-hiPSC-CMs

The effect of DTA-treatment on expression of proteins involved in Ca^2+^ handling prompted us to investigate intracellular Ca^2+^ release and dynamics in spontaneously beating DTA-treated and untreated hiPSC-PMs. To this aim, we employed non-ratiometric Fluo4-AM Ca^2+^-sensitive dye at day 40 of differentiation (Fig 3). To investigate SR Ca^2+^ handling, we analyzed both, spontaneous Ca^2+^ transients and the SR Ca^2+^ load induced by 10mM caffeine. The rate of spontaneous Ca^2+^ transients was significantly increased in DTA-treated PM-hiPSC-CMs compared to untreated group (Fig 3B). No significant difference was observed in Ca^2+^ transient amplitude (Ca_T_) between the two groups (Fig 3B). However, the caffeine-induced Ca^2+^ transient amplitude (Ca_SR_) was significantly increased in DTA-treated PM-hiPSC-CMs compared to untreated ones (Fig 3B). Consequently, the SR Ca^2+^ fractional release (SR FR), obtained as the ratio between Ca_T_ and Ca_SR_, was significantly reduced in DTA-treated compared to untreated PM-hiPSC-CMs (Fig 3B). The Ca_SR_ decay time (τ decay) was significantly reduced in DTA-treated compared to untreated PM-hiPSC-CMs (Fig 3B), suggesting an improvement in the SERCA2a Ca^2+^-reuptake in the SR. These data show that DTA-treatment protocol improves PM-hiPSC-CMs Ca^2+^- handling and are consistent with proteomics analysis indicating up-regulation of SR Ca^2+^-related proteins. Taken together, these data validate DTA as valuable treatment to improve functional maturation of Ca^2+^ handling in PM-hiPSC-CMs. Consequently, we employed DTA treatment to differentiate PM-hiPSC-CMs and perform further functional and electrophysiological characterization of PM-hiPSC-CMs at day 40.

**Fig 3.**
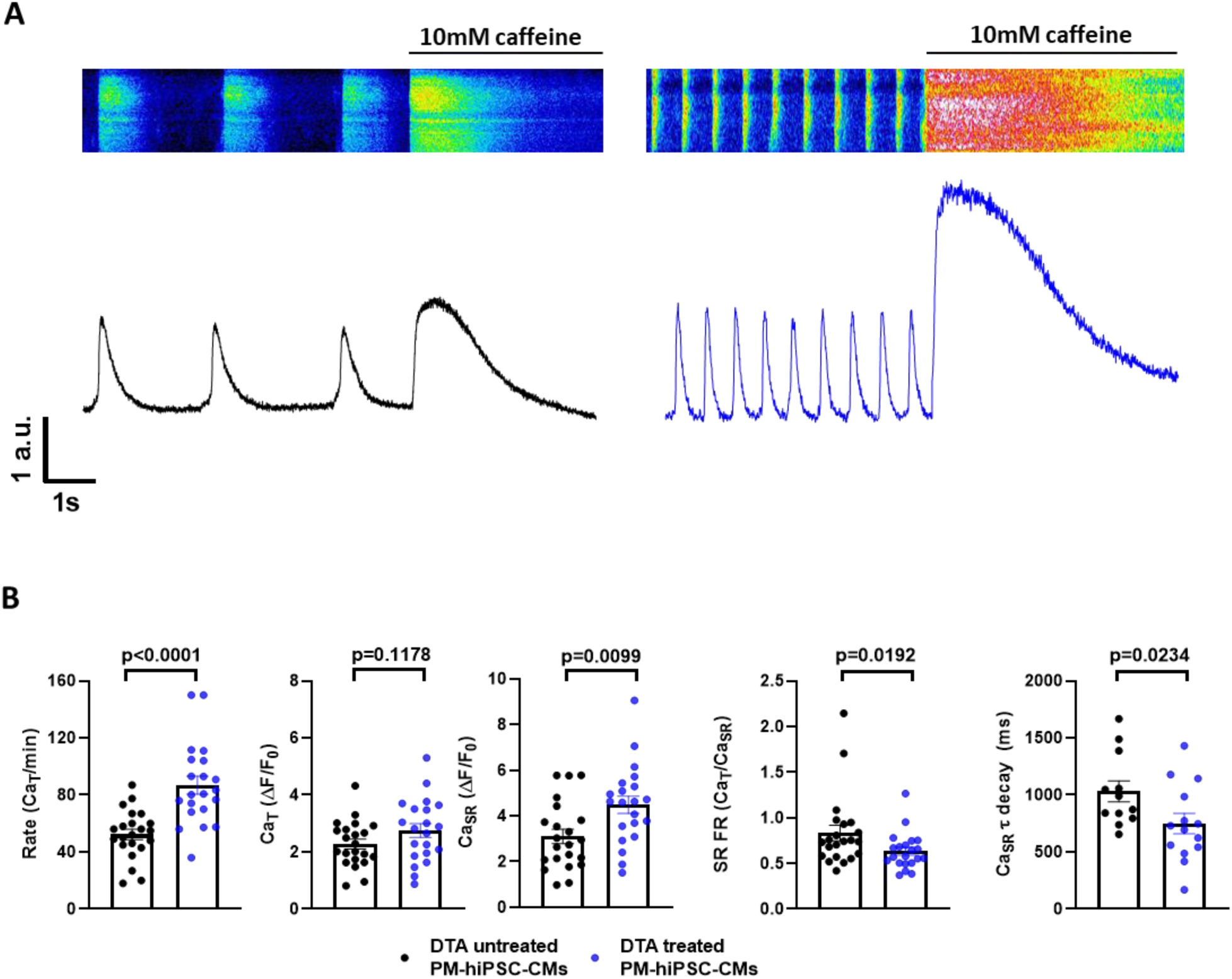
DTA treatment promotes Ca^2+^ handling maturation of PM-hiPSC-CMs. (A) Confocal representative traces (above) and averaged traces (bottom) of Ca^2+^ fluorescence generated by Fluo4-AM (5µM) in DTA-treated and untreated PM-hiPSC-CMs at day 40 of differentiation exposed to Tyrode and then to saturating concentration of caffeine (Caff, 10 mM). (B) Averaged rate of Ca^2+^ transients, amplitudes of Ca^2+^ transients (Ca_T_) in Tyrode and after superfusion of Caff (Ca_SR_), ratio between Ca_T_ and Ca_SR_ (fractional release, FR), τ decay of Ca_SR_ in DTA-treated PM-hiPSC-CMs compared to untreated group at day 40 post-differentiation. DTA-treated PM-hiPSC-CMs (d=5, n ≥ 12), DTA-untreated PM-hiPSC-CMs (d=4, n *≥* 13). Statistic: unpaired non-parametric Kruskal-Wallis test followed by Dunn’s multiple comparison test.

### DTA-treated PM-hiPSC-CMs display typical ionic currents of native SAN cells

Rhythmic generation of action potentials (APs) in SAN myocytes is contingent on diastolic depolarization (DD), which drives the membrane potential from the end of the repolarization phase of an AP to the threshold of the following AP. Key ionic currents involved in DD are the hyperpolarization-activated “funny”, cyclic nucleotide-gated inward current (I_f_,) ^42^, encoded by f-(HCN1-4) channels, the T-type current (I_CaT_), encoded by T-type Ca_v_3.1 channel ^43^ and the L-type Ca^2+^ current (I_CaL_) ^30,44,45^, encoded by L-type Ca_v_1.2 and Ca_v_1.3 channels ^30,43^. Although the AP upstroke of pacemaker cells is mainly driven by Ca^2+^ rather than Na^+^, two different Na^+^ current (I_Na_) components have been reported in the SAN: a TTX-sensitive “neuronal” I_Na_ pre-dominantly encoded by the Na_V_1.1 and Na_V_1.3 isoforms, and TTX-resistant I_Na_, encoded by the Na_V_1.5 isoform^46,47^.

Electrophysiological recordings were performed to investigate if DTA-treated PM-hiPSC-CMs presented ionic currents typical of pacemaker phenotype (Fig 4). Consistent with typical native SAN cells, I_f_ activated in a voltage-dependent manner upon membrane hyperpolarization in DTA-treated PM-hiPSC-CMs (Fig 4A; I_f_ half-activation voltage (V_0.5_) calculated from the steady-state activation curves is shown in inset of Fig 4A. 2 mM Cs^+^ almost completely blocked I_f_ in DTA-treated PM-hiPSC-CMs (Fig 4B). Recordings of Ca^2+^ currents were performed distinguishing I_CaT_ and I_CaL_ in DTA-treated PM-hiPSC-CMs (Figs 4C-D). First, total Ca^2+^ current (I_CaTOT_= I_CaL_ + I_CaT_) current-to-voltage (I-V) relationship was measured by applying depolarizing steps in the range of -80 to +60 mV from a HP of -80 mV (Fig 4C). I_CaL_ I-V relationship was obtained by applying depolarizing steps in the range of -55 to +60 mV from a HP of -55 mV (Fig 4C). Finally, I_CaT_ I-V relationship was calculated by subtracting the I_CaL_ I-V relationship from I_CaTOT_ (Fig 4C). Consistent with data from native SAN ^30,44,45^, maximum I_CaT_ peak was negatively shifted compared to that of I_CaL_ in DTA-treated PM-hiPSC-CMs (-20 mV for I_CaT_ vs -10 mV for I_CaL_ respectively) (Fig 4C). In addition, 3µM of the L-type dihydropyridine (DHP) blocker nifedipine (Nife) was used to evaluate block of I_CaL_ by applying a single step protocol (from -55 mV to -10 mV) (Fig 4D). 3µM Nife almost completely blocked I_CaL_ in DTA-treated PM-hiPSC-CMs (Fig 4D). Residual Nife-insensitive current suggested that a fraction of total population of L-type Ca_v_1.3 channels was not blocked by the DHP at steady-state.

**Fig 4.**
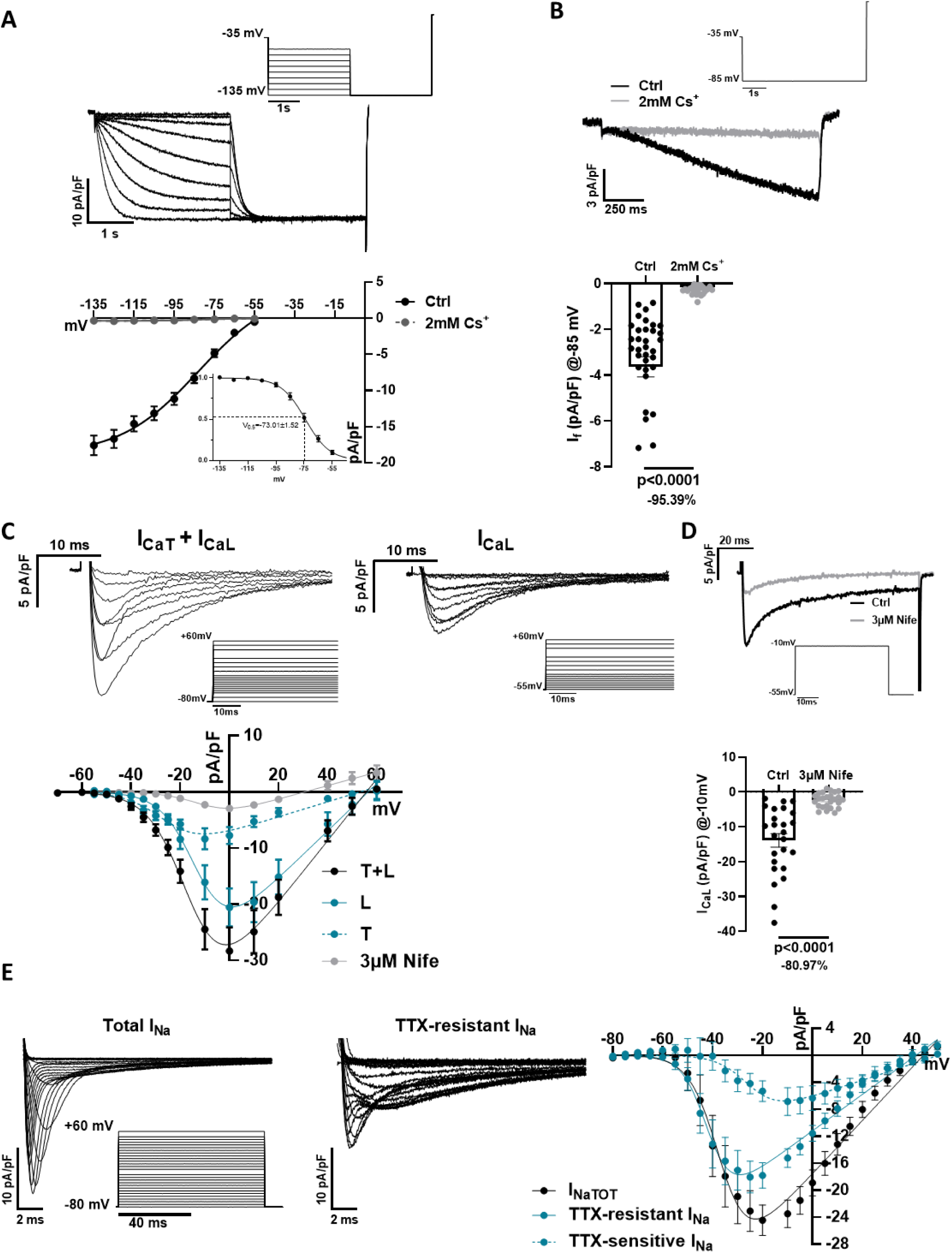
Ion channels characterization of DTA-treated PM-hiPSC-CMs. (A) Voltage-clamp (V-clamp) protocol used for I_f_ recordings and representative traces of I_f_ (above). Averaged current-to-voltage (I-V) I_f_ relationship measured in Tyrode (Ctrl) and after 2mM Cs^+^ superfusion in dissociated DTA-treated PM-hiPSC-CMs at day 40 of differentiation (d=4, n≥23) (bottom). Half-activation voltage (V_0.5_) calculated from steady-state activation curve (inset) in dissociated DTA-treated PM-hiPSC-CMs at day 40 of differentiation (d=4, n=28). (B) I_f_ was activated by -85mV hyperpolarizing step protocol and inhibited by 2mM Cs^+^ (above). Histogram representing averaged I_f_ density in control (Ctrl) and after 2 mM Cs^+^ superfusion in dissociated DTA-treated PM-hiPSC-CMs at day 40 of differentiation (d=7, n=36) (bottom). Statistic: paired non-parametric Wilcoxon test. (C) V-clamp protocol from a holding potential (HP) of − 80 mV used for I_CaT+L_ sample trace recording (left) and V-clamp protocol from a HP of − 55 mV used for I_CaL_ sample trace recording (right) in DTA-treated PM-hiPSC-CMs (above). I-V relationship of Ca^2+^ current recorded from a HP = − 80 mV (black circles) or from HP = − 55 mV (blue circles) measured in dissociated DTA-treated PM-hiPSC-CMs at day 40 post-differentiation (d=6, n=20). The dashed line indicates the net I_CaT_ I–V curve, calculated as the difference between values obtained from HP of − 55 mV from those from HP − 80 mV. I-V relationship of Ca^2+^ current was measured after superfusion of 3µM Nifedipine (Nife) in dissociated DTA-treated PM-hiPSC-CMs at day 40 post-differentiation (d=3, n=10). (D) I_CaL_ was activated by -10mV hyperpolarizing step protocol and inhibited by Nife (above). Histogram representing averaged I_CaL_ density in Ctrl and after Nife superfusion in dissociated DTA-treated PM-hiPSC-CMs at day 40 post-differentiation (d=6, n=25) (bottom). Statistic: paired non-parametric Wilcoxon test. (E) Representative traces of total I_Na_, TTX-resistant I_Na_ and V-clamp protocol used for recordings (inset). Averaged total I_Na_, TTX-resistant and TTX-sensitive I_Na_ I-V relationships measured in dissociated DTA-treated PM-hiPSC-CMs at day 40 of differentiation (d=3, n=10).

Total I_Na_ (I_NaTOT_ = TTX-resistant I_Na_ + TTX-sensitive I_Na_) I-V relationship was measured by applying depolarizing steps to the range of -80/+60 mV from a HP of -80 mV (Fig 4E). TTX-resistant I_Na_ I-V relationship was obtained by applying 100nM TTX (Fig 4E). Finally, TTX-sensitive I_Na_ I-V relationship was calculated by subtracting the TTX-resistant I_Na_ I-V relationship from I_NaTOT_ (Fig 4E). Both TTX-resistant and TTX-sensitive I_Na_ were detected in DTA-treated PM-hiPSC-CMs (Fig 4E).

### Chronotropic regulation of spontaneous activity of DTA-treated PM-hiPSC-CMs by autonomic agonists

DD generates the automaticity of the native SAN. APs obtained from DTA-treated PM-hiPSC-CMs were recorded (Fig 5). At baseline, spontaneous AP rate was 163 ± 1 bpm with a maximum diastolic potential (MDP) of -53.6 ± 1.7 mV and an amplitude of 92.6 ± 3.1 mV in DTA-treated PM-hiPSC-CMs (Fig 5A). The maximum upstroke velocity (dV/dt_max_) was 46.7 ± 13 mV/ms in DTA-treated PM-hiPSC-CMs (Fig 5A). AP duration measured at 90% repolarization (APD_90_) was 141.4 ± 9.3 ms in DTA-treated PM-hiPSC-CMs (Fig 5A). APD_20_ and APD_50_ were also measured (Fig 5A).

**Fig 5.**
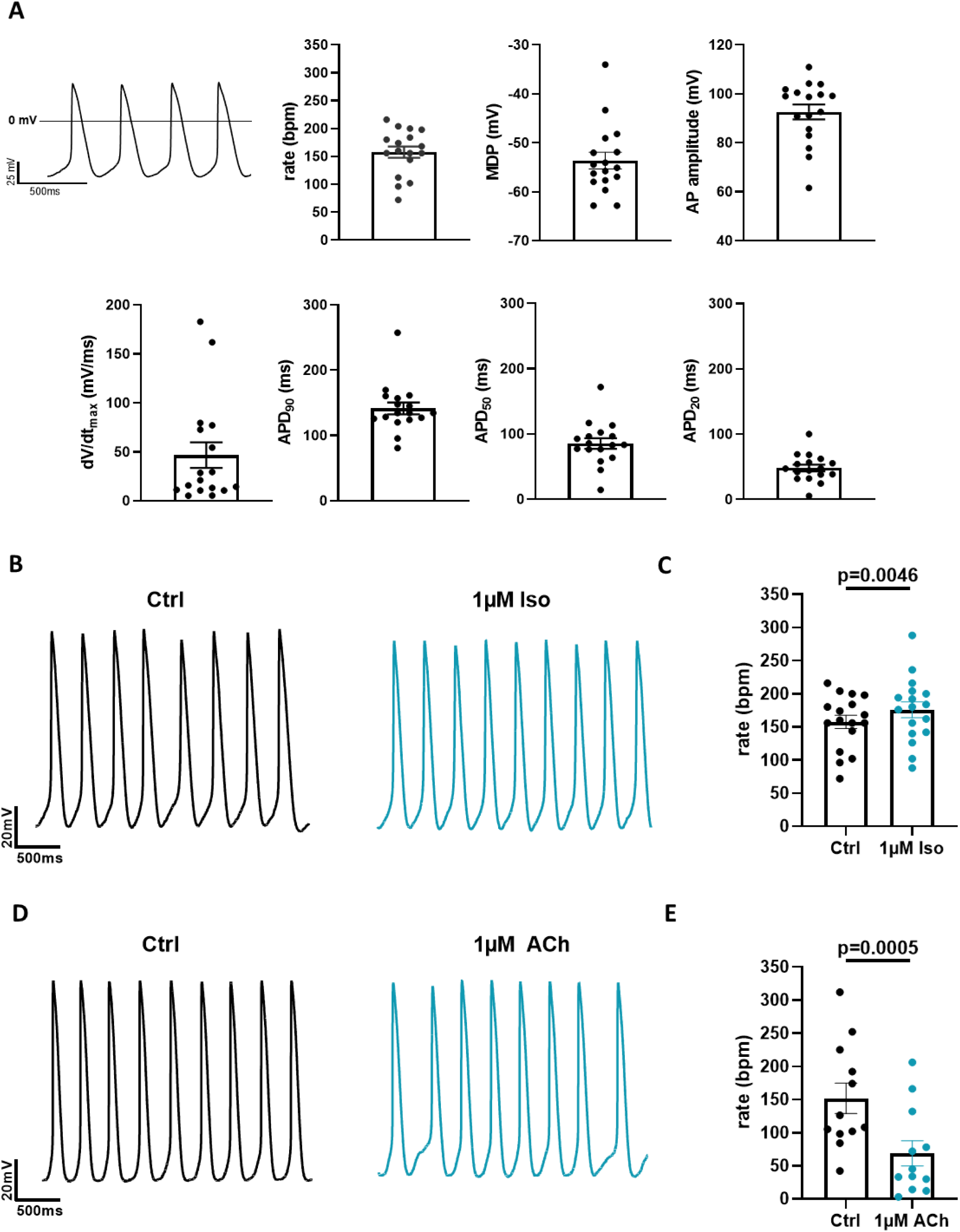
Modulation of automaticity in DTA-treated PM-hiPSC-CMs by autonomic agonists. (A) Examples of action potentials (APs) recorded using patch-clamp technique in gap-free mode and average parameters of APs (rate, minimum diastolic potential - E_diast_, AP amplitude, voltage on time ratio - dV/dt_max_, AP duration – APD in dissociated DTA-treated PM-hiPSC-CMs at day 40 after differentiation (d=3, n=17). (B) Representative recordings of APs in Ctrl and after superfusion of 1µM Isoprenaline (Iso) and (C) average AP rate before and after 1µM Iso superfusion in dissociated DTA-treated PM-hiPSC-CMs at day 40 of differentiation (d=3, n=17). Statistic: paired non-parametric Wilcoxon test. (D) Representative recordings of APs in Ctrl and after superfusion of 1µM Acetylcholine (ACh) and (E) average AP rate before and after 1µM ACh superfusion in dissociated DTA-treated PM-hiPSC-CMs at day 40 of differentiation (d=4, n=12). Statistic: paired non-parametric Wilcoxon test.

Activation of β-AR induces positive chronotropic response of SAN automaticity. Conversely, cholinergic agonists induce potent negative chronotropic effect on SAN automaticity ^48,49^. Like native SAN pacemaker cells, DTA-treated PM-hiPSC-CMs showed positive chronotropic response to β-adrenergic stimulation by 1µM Isoproterenol (Iso) (Figs 5B-C), resulting in significant increase in beating rates (Fig 5C). Stimulation of M2R by 1µM Acetylcholine (ACh) (Fig 5D-E) induced significant decrease in beating rate (Fig 5E).

### Modulation of ionic currents in DTA-treated PM-hiPSC-CMs by autonomic agonists

I_f_ ^50,51^ and Ca_v_1.3-mediated I_CaL_ ^28,30^ constitute important mechanisms in regulation of native SAN rate. Activation of β-ARs positively shifts the I_f_ activation curve, thereby supplying more inward current during the DD ^48^. At the same time, β-ARs stimulation induces an increase in the I_CaL_ ^30^. By contrast, activation of M2R by ACh negatively shifts the I_f_ activation curve decreasing the inward current during the DD ^48^ and diminishes the I_CaL_ ^51^. Consistently with these studies, I_f_ was significantly increased (Fig 6A) and decreased (Fig 6B) after Iso 1µM and 1µM ACh superfusion, respectively, at -85 mV in DTA-treated PM-hiPSC-CMs. In addition, I_CaL_ peak density was significantly increased by Iso (Fig 6C) and decreased (Fig 6D) after Iso 1µM and 1µM ACh superfusion, respectively, at -85 mV in DTA-treated PM-hiPSC-CMs. The G protein activated K^+^ current (I_KACh_) is required for the negative chronotropic response of native SAN pacemaker activity to cholinergic agonists ^52^. We recorded I_KACh_ by holding the membrane voltage at -45 mV and then perfusing 1µM ACh for 10 s before returning to control solution in DTA-treated PM-hiPSC-CMs (Fig 6E). Exposing DTA-treated PM-hiPSC-CMs to 1µM Ach, I_KACh_ was significantly evoked (Fig 6E). After the peak, *I_KACh_* showed typical decay kinetics as a result of desensitization (Fig 6E) ^53^.

**Fig 6.**
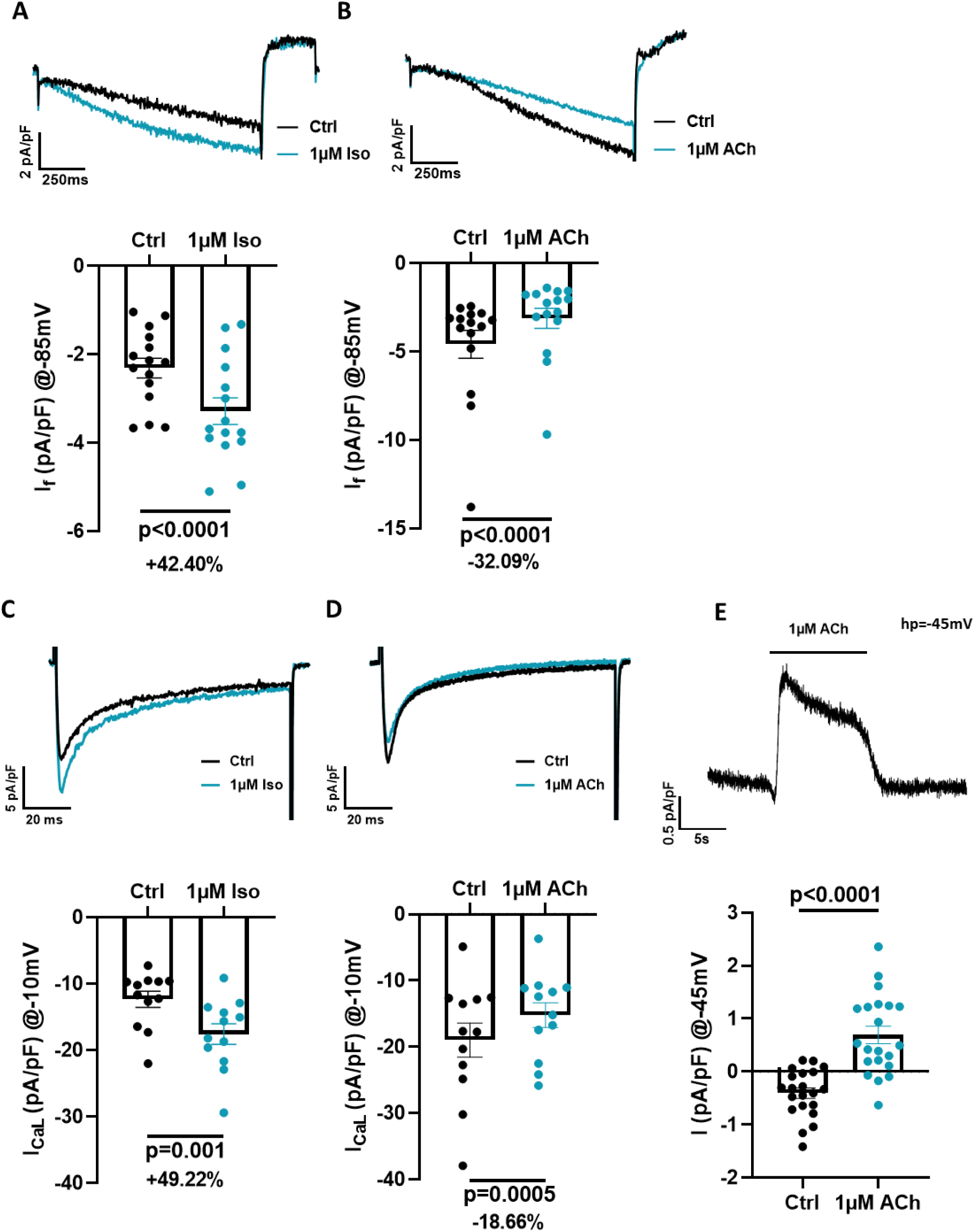
Regulation of ion channels in DTA-treated PM-hiPSC-CMs by autonomic agonists. (A) I_f_ was activated by -85mV hyperpolarizing step protocol and recorded in Ctrl and under superfusion of 1µM Isoprenaline (Iso). Representative traces of I_f_ in Ctrl and under superfusion of 1µM Iso (above). Averaged current densities of I_f_ before and after 1µM Iso superfusion in dissociated DTA-treated PM-hiPSC-CMs at day 40 after differentiation (d=3, n=15) (bottom). Statistic: paired non-parametric Wilcoxon test. (B) I_f_ was activated by -85mV hyperpolarizing step protocol and recorded in Ctrl and under superfusion of 1µM Acetylcholine (ACh). Representative traces of I_f_ in Ctrl and under superfusion of ACh (above). Averaged current densities of I_f_ before and after 1µM ACh superfusion in dissociated DTA-treated PM-hiPSC-CMs at day 40 after differentiation (d=3, n=15) (bottom). Statistic: paired non-parametric Wilcoxon test. (C) I_CaL_ was activated by -10mV hyperpolarizing step protocol and recorded in Ctrl and under superfusion of 1µM Iso. Representative traces of I_CaL_ in Ctrl and under superfusion of 1µM Iso (above). Averaged current densities of I_CaL_ before and after 1µM Iso superfusion in dissociated DTA-treated PM-hiPSC-CMs at day 40 after differentiation (d=3, n=12) (bottom). Statistic: paired non-parametric Wilcoxon test. (D) I_CaL_ was activated by - 10mV hyperpolarizing step protocol and recorded in Ctrl and under superfusion of 1µM ACh. Representative traces of I_CaL_ in Ctrl and under superfusion of 1µM ACh (above). Averaged current densities of I_CaL_ before and after 1µM ACh superfusion in dissociated DTA-treated PM-hiPSC-CMs at day 40 after differentiation (d=3, n=12) (bottom). Statistic: paired non-parametric Wilcoxon test. (E) Representative trace of 1µM ACh-sensitive endogenous current at HP=-45mV (above). Averaged current densities in Ctrl and under superfusion of 1µM Ach in dissociated DTA-treated PM-hiPSC-CMs at day 40 after differentiation (d=3, n=21) (bottom). Statistic: paired non-parametric Wilcoxon test.

## DISCUSSION

### Improving sinoatrial-like hiPSCs differentiation protocol

The use of hiPSC technology enabled mechanistic investigation of inherited pathologies of cardiac function using cardiomyocytes recapitulating the molecular and cellular environment of donor patients. Most hiPSC differentiation protocols have focused on obtaining ventricular-like hiPSC-CMs to model disorders affecting heart ventricular function^54,55^, while few others has proven useful to enrich hiPSC cultures in atrial-like CMs, particularly by using small molecules and retinoic acid (RA) in accurate timing and concentration ^56^.

To fate, five protocols have been proposed for differentiating hiPSC towards PM-hiPSC-CMs and are contingent on 2D and 3D matrix structures. Yechikov et al. ^14^ have shown that inhibiting the NODAL signaling pathway causes PITX2c downregulation in the cardiac mesoderm and greatly improves differentiation of hiPSC in PM-hiPSC-CMs ^14^. Protze et al. ^13^ and Liu et al. ^12^ reported that the fraction of PM-hiPSC-CM present in hiPSC-CMs population can be significantly enriched by simultaneous manipulation of BMP, FGF, and RA signaling pathways. Schweizer and coworkers ^15^ using a culture media-based approach, reported that co-culturing hiPSC with visceral endoderm-like cells (END2 cells) in a serum-free medium, promotes their differentiation into spontaneously beating clusters of PM-hiPSC-CMs having characteristics of SAN-like cardiomyocyte. The same authors have recently proposed a new protocol to differentiate PM-hiPSC-CMs based on sequential modulation of the Wnt pathway ^57^. Here, we have modified the protocol reported by Liu et al. ^12^, adapting the 2D matrix-sandwich method which promotes epithelial-to-mesenchymal transition, a critical step for generating cardiomyocytes^21^. Current investigations focus on feeding ventricular-like hiPSC-CMs with fatty acids in order to activate the mitochondrial oxidative phosphorylation and improve their maturity ^58,59^. Conversely, SAN-like cardiomyocytes are known to exhibit a different metabolic profile characterized by lower oxygen consumption, high activity of glycolytic pathway and weaker contractile properties than ventricular cardiomyocytes^32^. Consequently, in the present study, we differentiated the hiPSCs in glucose-rich, low fatty acid containing RPMI 1640+B27 medium to promote glycolytic metabolism than the oxidative one.

In addition, we have functionally improved the PM-hiPSC-CM population by adding Dex, T3 and cAMP (‘DTA’ treatment) at specific time window (from day 32 till day 40) of the differentiation protocol to improve Ca^2+^-handling (Dex and T3) and conduction (cAMP).

### DTA cocktail affects the calcium handling machinery

Recent studies have indicated that Dex and T3 treatment improves the key properties of the hiPSC-CMs by improving intracellular calcium handling machinery ^16–18^. Since Dex accelerates Ca^2+^ transient decay enhancing cardiac SR Ca^2+^ ATPase (SERCA2a) and Na^+^-Ca^2+^exchanger (NCX1) function in human embryonic stem cell derived cardiomyocytes (hESC-CMs) ^19^, it is considered a good candidate molecule to promote hiPSC-CM maturation when used in combination with other factors such as T3. Especially, it was shown that T3 and Dex together enhance the functional coupling between L-type Ca^2+^ channels and RyR2 on SR in hiPSC-CMs, ameliorating the Ca^2+^-induced Ca^2+^-release mechanism^17^. Furthermore, persistent activation of the cAMP pathway contributes to hiPSC-CM maturation, by positively regulating the assembly of gap junctions ^16^.

In our study, we show that our DTA-based protocol drives hiPSC differentiation towards cell population showing typical functional hallmarks of native pacemaker myocytes (Figs 1B-C). Differences in terms of maturation from day -1 are more pronounced in DTA-treated than in untreated PM-hiPSC-CMs (Fig 1B). As expected, DTA treatment upregulated expression of proteins regulating intracellular Ca^2+^ handling (Figs 1B-C). It is noteworthy that besides the activity of several ion channels, evidence abounds that the activity of SR via RyR2-dependent Ca^2+^ release is also involved in SAN pacemaking ^60–63^. Indeed, it has been proposed that the crosstalk between RyR2-dependent rhythmic Ca^2+^ release from the SR and sarcolemmal channels such as I_f_ and Ca_v_1.3-mediated I_CaL_ generates SAN automaticity ^28,61,63,64^. In comparison to untreated PM-hiPSC-CMs, DTA-treated PM-hiPSC-CMs displayed increased levels of protein involved in regulation of Ca^2+^ handling (*i.e.*, RYR2 and CAMK2D) at day 40 of differentiation (Fig 1C). Moreover, after having reached a peak during cardiac mesoderm stage, expression of typical markers of ventricular differentiation, HAND1 and NKX2-5, were reduced in DTA-treated PM-hiPSC-CMs at day 40 of differentiation (Fig 1D).

Consistent with these observations, GO analysis (Fig 2) of DTA-treated PM-hiPSC-CM showed up-regulation of key pathways underlying cardiac conduction^60,65^, intracellular calcium homeostasis^60–62^ and sodium ion transport regulation^46,47^ , previously showed to be involved in the pacemaker mechanism. We also found significantly enriched pathways in DTA-treated PM-hiPSC-CM related to cellular respiration/metabolism^32^, two important and interconnected biological processes in native pacemaker cells. Indeed, Senges and coworkers^33^ reported that hypoxia depressed automaticity and conduction in rabbit SAN demonstrating the importance of oxygen levels for cardiac pacemaking. Furthermore, it has been shown that native pacemaker myocytes are endowed with a complex mitochondrial network crucial for cell survival and for supplying energy by oxidative phosphorylation process^34^. The essential role of normal-functioning mitochondrial-linked processes in pacemaker cells is also highlighted by data showing that mitochondrial dysfunction reduces SAN electrical activity^66^. GO analysis also uncovered up-regulation of differential expressed proteins related to the regulation of iron ion transport. Iron plays a key role in physiological and intracellular functions in myocytes and several studies have showed detrimental effects of iron overload or deficiency^67,68^. Further reinforcing the concept that DTA treatment drives hiPSC-CM differentiation towards a more specific pacemaker cell phenotype, functional GO enrichment analysis unmasked multiple biological upregulated processes in untreated PM-hiPSC-CM which are not linked to cardiomyocyte development and, particularly, with cardiac automaticity. Consistently with the limited proliferative capacity of cardiac cells^69,70^, DTA-treated PM-hiPSC-CM showed down-regulation of pathways involved in cell cycle progression and cell proliferation in comparison to DTA-untreated PM-hiPSC-CM. The vascular endothelial growth factor (VEGF) signaling pathway was also significantly expressed in DTA-untreated PM-hiPSC-CM. Intriguingly, it has been showed that elevated levels of VGEF can acutely increase the susceptibility to atrial arrythmias, in particular atrial fibrillation^71^. These data may thus suggest that cells treated with DTA, in which VEGF pathway is downregulated present with more stable pacemaker activity. Epithelial to mesenchymal transition (EMT) pathway is upregulated in untreated PM-hiPSC-CMs. In cardiac development, the EMT pathway underlies differentiation of a new population of cells, referred to as Epicardially Derived Cells (EPDCs). Migration of EPDCs into the myocardium, contributes to the formation of coronary endothelial cells, smooth muscle cells, cardiac fibroblasts and play an important role in the induction of the ventricular compact myocardium and the differentiation of the Purkinje fibers^72^. Thus, downregulation in pathways related to EMT may reinforce pacemaker identity of DTA-treated PM-hiPSC-CMs. Furthermore, GO analysis revealed up-regulation in untreated cells of DEPs involved in the regulation of stress-fibers assembly, and particularly of positive regulation of actin filament bundle assembly, a biological process fundamental for cardiomyocytes contractility but not prominently involved in cardiac automaticity^73–75^.

At functional level, DTA-treated PM-hiPSC-CMs showed faster Ca^2+^ transient rate of spontaneous Ca^2+^ transients in comparison to the untreated PM-hiPSC-CMs (Fig 3). Ca_SR_ was increased after caffeine pulse and SR Ca^2+^ re-uptake was faster in DTA-treated PM-hiPSC-CMs compared to untreated group (Fig 3B). Consequently, SR FR was clearly decreased in DTA-treated PM-hiPSC-CMs (Fig 3C), which is indicative of increased efficiency of Ca2+ release from SR, in response to caffeine stimulus.

In sum, DTA treatment of PM-hiPSC-CMs ameliorates the differentiation process, in particular in Ca^2+^ handling dynamic, as reported by protein expression/GO and functional data (Fig 1-3). Finally, up-regulation of SAN markers (Fig 1B-C) accompanied with down-regulation of ventricular markers (Fig 1D) is strongly indicative of a differentiation of hiPSCs to a more SAN- rather than ventricular-like phenotype.

### DTA-treated PM-hiPSC-CMs recapitulate native SAN features

In this study, we report that DTA-treated PM-hiPSC-CMs at day 40 of differentiation express ionic currents underlying SAN pacemaker activity: I_f_, I_CaL,_ I_CaT_ and I_Na_ (Fig 4). In addition, we recorded robust automatic action potentials (Fig 5A). Automaticity was faster than the intrinsic heart rates recorded in humans at rest, but was similar to heart rates recorded in 24 hours term-born infants ^76^. Automaticity of DTA-treated PM-hiPSC-CMs was regulated in opposite directions by the sympathomimetic isoproterenol and acetylcholine, similarly to native SAN pacemaker activity. Importantly, in DTA-treated PM-hiPSC-CMs, activation of β-ARs by isoproterenol upregulated two key ionic currents involved in the chronotropic response, I_f_ ^50,51^ and I_CaL_^30,51^ (Figs 5C-D and Figs 6A-C). In line with the hypothesis that PM-hiPSC-CMs recapitulate the chronotropic mechanism of native SAN myocytes, I_f_ and I_CaL_ were negatively regulated by the activation of M2Rs (Figs 5E-F and Figs 6B-D). I_KACh_ is required for regulation of native SAN pacemaker activity and is an important downstream effector of the parasympathetic nervous system ^52^. DTA-treated PM-hiPSC-CMs expressed *I_KAch_* presenting typical hallmarks of native SAN *I_KAch_*, namely robust activation upon Ach perfusion and time-dependent decay of current as the result of channel desensitization (Fig 6E) ^53^.

In conclusion, our study indicate that DTA treatment applied to Liu et al. ^12^ differentiation protocol supports maturation of PM-hiPSC-CMs, by ameliorating Ca^2+^ handling, improving the expression of key SAN markers and upregulating cellular processes linked to SAN automaticity. This bulk of evidences supports the view that DTA-treated hiPSC-CMs constitute a faithful model of SAN myocytes *in vitro*. The ability to generate functional SAN myocytes from hiPSCs provides an unprecedented opportunity to study human pacemaker development and function, to model diseases that affect this subpopulation of cardiomyocytes and to design cell-based therapies for patients with SAN dysfunction.

